# Molecular Mimicry Between Epstein-Barr Virus and Human Herpesvirus-6 Proteins and Central Nervous System Proteins: Implications for T and B Cell Immunogenicity in an In Silico Study

**DOI:** 10.1101/2025.04.02.646883

**Authors:** Abbas F. Almulla, Muslimbek G. Normatov, Thitiporn Supasitthumrong, Michael Maes

## Abstract

**Background:** The Epstein-Barr virus (EBV) and human herpesvirus 6 (HHV-6) are frequently linked to neuropsychiatric illnesses such as multiple sclerosis, depression, and chronic fatigue syndrome/myalgic encephalomyelitis. These viruses may induce autoimmune reactions by molecular mimicry, leading to damage to self-epitopes in the central nervous system (CNS).

**Objective:** This study seeks to explore the common pentapeptides present in EBV and HHV-6 viral antigens alongside various CNS-related proteins via molecular mimicry. Additionally, it will assess the immunogenicity of these shared pentapeptides in T and B cells.

**Method:** Sequence alignment was conducted to assess molecular mimicry between 32 EBV and HHV-6 antigens and 10 CNS autoantigens. Protein sequences were obtained from UniProt, structural homology was analyzed using AlphaFold and PyMol, and shared pentapeptides were identified with Alignmentaj. Immunogenicity was assessed via the Immune Epitope Database (IEDB) for potential T- and B-cell activation.

**Results:** A total of 91 mimicry pentapeptides were identified between viral antigens (EBV and human HHV-6), and CNS proteins. Notably, synapsin (SYN)1 exhibited the highest mimicry, sharing multiple pentapeptides with EBV nuclear antigen (EBNA)1, EBNA6, latent membrane protein (LMP)1, and early antigen diffused (EA-D) and 6 different HHV-6 antigens. Myelin proteins including myelin basic protein, myelin-associated glycoprotein, and myelin-oligodendrocyte glycoprotein also displayed shared pentapeptides with EBV/HHV-6 antigens, indicating potential immune cross-reactivity. EBNA1, EBNA2, EBNA6, LMP1, LMP2, EA-D, and BLLF1 structurally resemble CNS autoantigens and act as immunoreactive epitopes for human T and B cells. Except for EBNA2 and protein U94, all share immunogenic pentapeptide sequences with SYN1.

**Conclusion:** EBV and HHV-6 antigens mimic CNS proteins, potentially triggering autoimmune responses via T and B cell activation. Shared pentapeptides suggest a link between viral infections and CNS autoimmunity. Further research is needed to clarify molecular mechanisms and explore targeted therapies to mitigate virus-induced neuroinflammation.

## Introduction

The phenomenon whereby foreign antigens, such as those derived from pathogens, exhibit similar structures to self-antigens is known as molecular mimicry, which is implicated in triggering autoimmune responses [1, 2]. The pathogenesis of several major autoimmune diseases, including those related to the central nervous system (CNS), involves this phenomenon [1, 3, 4]. Structurally similar antigens, when presenting to the immune system, are able to generate immune responses mistakenly targeting host tissues, leading to damage as a result of the inability of the immune system to distinguish self from non-self-antigens [2, 5, 6]. For example, rheumatism exemplifies an immune response that simultaneously affects cardiac tissues due to streptococcal antigens [6]. In CNS diseases such as multiple sclerosis (MS), where the autoreactive T cells are known to target myelin proteins, molecular mimicry was frequently reported as a key mechanism [7].

Pathogens such as viruses and bacteria contain antigens that exhibit structural similarities to myelin proteins, resulting in cross-reactive immune responses [6, 8]. A growing body of evidence linked infections caused by viruses like Epstein-Barr virus (EBV), human herpesvirus 6 (HHV-6), Severe acute respiratory syndrome coronavirus 2 (SARS CoV 2), and Zika virus (ZikV) with autoimmune CNS diseases, including but not limited to MS [9–11]. França et al. revealed an association between infection with ZikV and neurological complications, as its NS5 protein exhibits homology with CNS antigens, triggering autoimmune responses through mimic the former antigens [9]. Additionally, SARS-CoV-2 has been linked to autoimmune diseases in genetically susceptible individuals, further underscoring the impact of viral infections on CNS pathology [10]. These findings underscore the viruses’ potential to trigger autoimmune demyelinating processes via molecular mimicry.

Recently, our findings suggested that the reactivation of EBV and HHV-6 is critical in maintaining autoimmune processes in MS patients [12]. These results demonstrated that autoantibodies against myelin proteins, such as myelin basic protein (MBP) and myelin oligodendrocyte glycoprotein (MOG), in those patients are associated with reactivation of EBV and HHV-6 [12]. Similarly, in Long COVID disease, our results revealed significant correlations between HHV-6 reactivation and autoantibodies (immunoglobulins A and G) targeting zonulin and occludin, which are tight junction proteins. HHV-6 and EBV reactivation and consequent autoimmune responses to CNS self-epitopes may play a role in Long COVID, depression, chronic fatigue syndrome/myalgic encephalomyelitis (CFS/ME) [13–16]. The findings emphasize that these viruses’ reactivation has key roles in triggering or sustaining autoimmune responses [14].

CNS proteins, including MBP, MOG, and proteolipid (PLP), are essential for maintaining myelin integrity and neural conduction [17, 18]. Disruption of these proteins through immune-mediated attacks results in demyelination and impaired neuronal function, as noted in MS [19]. Tight junction proteins such as claudin-5 and occludin, which maintain blood-brain barrier (BBB) integrity alongside synaptic protein such as synapsins, are also targeted in autoimmune CNS disorders, exacerbating inflammation and neurodegeneration [20–22]. It is important to note that molecular mimicry is not the sole mechanism in the pathophysiology of CNS diseases; other mechanisms, such as epitope spreading, bystander activation, and immune-inflammatory responses, are also involved [11, 23, 24]. Miller et al. reported that T cell responses targeting the immunodominant myelin PLP-139-151 epitope did not result from cross-reactivity between Theiler’s murine encephalomyelitis virus and self-epitopes, commonly referred to as molecular mimicry. Instead, these responses were attributed to de novo priming of self-reactive T cells to sequestered autoantigens released as a consequence of virus-specific T cell-mediated demyelination, known as epitope spreading [25].

EBV and HHV-6 exhibit neurotrophic properties that substantially influence CNS disease. These viruses establish latency in host cells, avoid immune surveillance, and periodically reactivate, presenting continuous threats to brain health [26, 27]. Lanz et al. revealed a molecular mimicry between EBV antigen (namely EBV transcription factor EBV nuclear antigen 1, EBNA1 and glial cell adhesion molecule (GlialCAM) in MS disease [28]. HHV-6 can impair cellular immunity by targeting CD4+ T cells, further complicating immune dysregulation [29]. Likewise, Mameli et al. reported that HHV-6 antigens mimic MBP resulting in autoimmune responses against the latter [30]. Moreover, EBV and HHV-6 viral proteins mimic important CNS proteins like α and β-synuclein, neurofilament light chain [31, 32], anoctamin 2 [33], α-B crystallin (CRYAb) [34] and Septin-9 [35], suggesting contributions of these mechanisms in the pathogenesis of neuroautoimmune diseases. Kanduc (2013) reported that pentapeptides can induce highly specific antibodies and mediate precise immune interactions [36]. However, our understanding of mimicry across a broader spectrum of CNS proteins is still not fully comprehended, as most research focuses on a limited range of proteins.

This study investigates the molecular mimicry of similar pentapeptides between 32 EBV and HHV-6 proteins and various CNS proteins, including myelin proteins (MBP, MOG, PLP), BBB proteins (occludin, claudin-5), cytoskeletal proteins (tubulin), and synaptic proteins (synapsin). We will examine whether the identified shared pentapeptides are immunogenic to T and B cells. The central hypothesis suggests that particular amino acid sequences in viral proteins structurally mimic CNS proteins, eliciting immune responses that may contribute to autoimmune and neuroinflammatory disorders, including MS.

## Material and methods

### Data retrieval

To examine molecular mimicry between herpesviruses (EBV and HHV-6) antigens and CNS autoantigens, 32 immunogenic viral antigens were selected. The EBV antigens include EBNA1, EBNA2, EBNA6 Latent Membrane Protein (LMP) 1, LMP2A/LMP2B, Early Antigen-Diffuse (EA-D), Early Antigen-Restricted (EA-R), Epstein-Barr Virus Glycoprotein gp350/220 (BLLF1), Glycoprotein B (gB), Glycoprotein H (gH), Major Capsid Protein (MCP), and Glycoprotein E (gE). The HHV-6 antigens include immediate-early protein 2 (U90/U87/U86), MCP, ribonucleotide reductase large subunit (RIR1), U24 protein, 120 kDa Glycoprotein O (U47), Large structural phosphoprotein (U11), DNA binding protein (DBP), putative CC-type chemokine (U83), DNA polymerase catalytic subunit (U38), capsid scaffolding protein (U53), large tegument protein deneddylase (U31), gH, glycoprotein (U21), Glycoprotein Q2 (U100), gB, G-protein coupled receptor homolog (U12), Glycoprotein 105 (U96/U97/U98/U99/U100), Uracil-DNA glycosylase (U81), G-protein coupled receptor homolog (U51), and U94 protein .

These viral proteins were examined for their structural homologies (shared pentapeptides) with 10 selected human CNS autoantigens (MBP, MOG, MOBP, MAG, SYN1, SYN2, CLDN5, OCLN, TUBA1A and TUBB), which serve as potential targets in the autoimmune neuropathies observed in MS, ME/CFS, and affective syndromes. The amino acid sequences of EBV and HHV-6 antigens and human CNS autoantigens were obtained from the Uniprot database [37] in fasta format. The ID number of EBV and HHV-6 antigens along with human CNS autoantigens in the UniProt database is shown in **Table 1**.

**Table 1.**
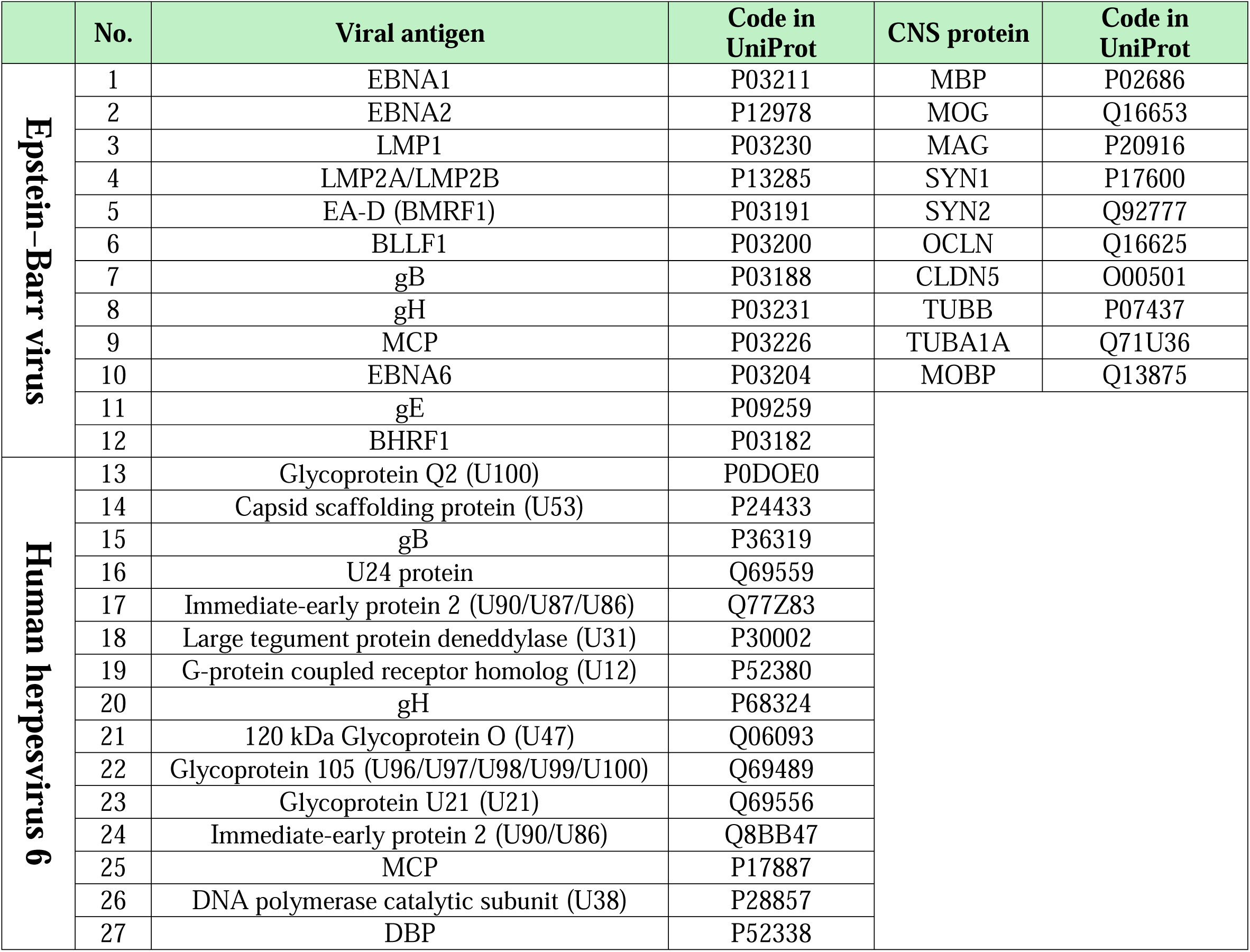
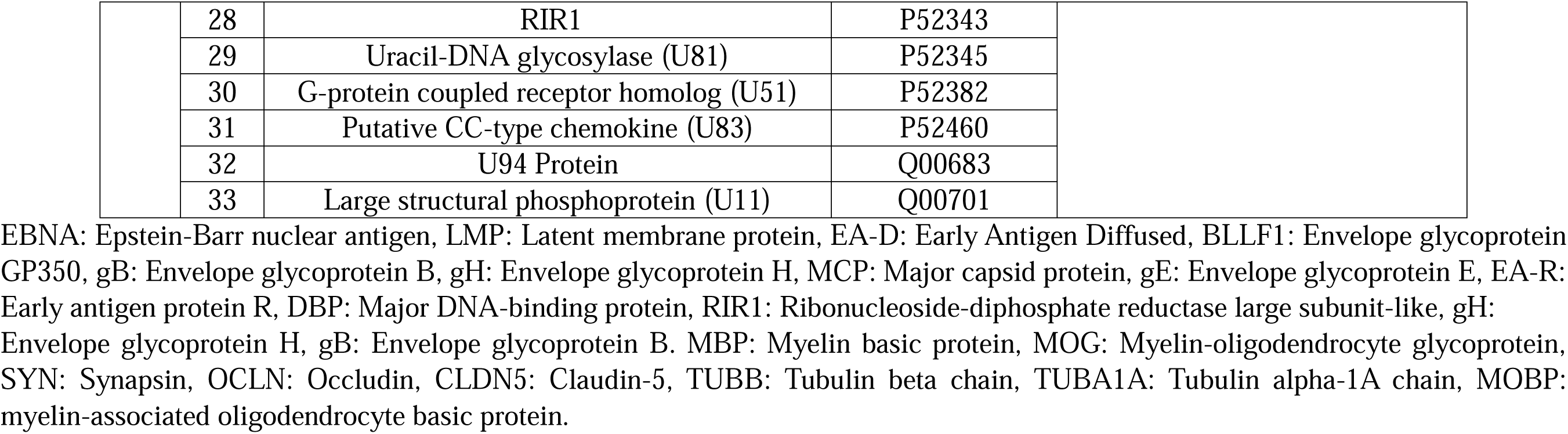
UniProt code for Epstein–Barr virus (EBV), Human herpesvirus 6 (HHV-6) and central nervous system (CNS) antigens utilized in the current study.

|  | No. | Viral antigen | Code in UniProt | CNS protein | Code in UniProt |
| --- | --- | --- | --- | --- | --- |
| **Epstein–Barr virus** | 1 | EBNA1 | P03211 | MBP | P02686 |
|  | 2 | EBNA2 | P12978 | MOG | Q16653 |
|  | 3 | LMP1 | P03230 | MAG | P20916 |
|  | 4 | LMP2A/LMP2B | P13285 | SYN1 | P17600 |
|  | 5 | EA-D (BMRF1) | P03191 | SYN2 | Q92777 |
|  | 6 | BLLF1 | P03200 | OCLN | Q16625 |
|  | 7 | gB | P03188 | CLDN5 | O00501 |
|  | 8 | gH | P03231 | TUBB | P07437 |
|  | 9 | MCP | P03226 | TUBA1A | Q71U36 |
|  | 10 | EBNA6 | P03204 | MOBP | Q13875 |
|  | 11 | gE | P09259 |  |  |
|  | 12 | BHRF1 | P03182 |  |  |
| **Human herpesvirus 6** | 13 | Glycoprotein Q2 (U100) | P0DOE0 |  |  |
|  | 14 | Capsid scaffolding protein (U53) | P24433 |  |  |
|  | 15 | gB | P36319 |  |  |
|  | 16 | U24 protein | Q69559 |  |  |
|  | 17 | Immediate-early protein 2 (U90/U87/U86) | Q77Z83 |  |  |
|  | 18 | Large tegument protein deneddylase (U31) | P30002 |  |  |
|  | 19 | G-protein coupled receptor homolog (U12) | P52380 |  |  |
|  | 20 | gH | P68324 |  |  |
|  | 21 | 120 kDa Glycoprotein O (U47) | Q06093 |  |  |
|  | 22 | Glycoprotein 105 (U96/U97/U98/U99/U100) | Q69489 |  |  |
|  | 23 | Glycoprotein U21 (U21) | Q69556 |  |  |
|  | 24 | Immediate-early protein 2 (U90/U86) | Q8BB47 |  |  |
|  | 25 | MCP | P17887 |  |  |
|  | 26 | DNA polymerase catalytic subunit (U38) | P28857 |  |  |
|  | 27 | DBP | P52338 |  |  |

|  |  |  |  |
| --- | --- | --- | --- |
|  | 28 | RIR1 | P52343 |
|  | 29 | Uracil-DNA glycosylase (U81) | P52345 |
|  | 30 | G-protein coupled receptor homolog (U51) | P52382 |
|  | 31 | Putative CC-type chemokine (U83) | P52460 |
|  | 32 | U94 Protein | Q00683 |
|  | 33 | Large structural phosphoprotein (U11) | Q00701 |

EBNA: Epstein-Barr nuclear antigen, LMP: Latent membrane protein, EA-D: Early Antigen Diffused, BLLF1: Envelope glycoprotein GP350, gB: Envelope glycoprotein B, gH: Envelope glycoprotein H, MCP: Major capsid protein, gE: Envelope glycoprotein E, EA-R: Early antigen protein R, DBP: Major DNA-binding protein, RIR1: Ribonucleoside-diphosphate reductase large subunit-like, gH: Envelope glycoprotein H, gB: Envelope glycoprotein B. MBP: Myelin basic protein, MOG: Myelin-oligodendrocyte glycoprotein, SYN: Synapsin, OCLN: Occludin, CLDN5: Claudin-5, TUBB: Tubulin beta chain, TUBA1A: Tubulin alpha-1A chain, MOBP: myelin-associated oligodendrocyte basic protein.

The AlphaFold database [38] and the PyMol program [39] (https://www.pymol.org/) were used to illustrate the location of mimicry pentapeptides in the 3D surface structures of human nervous system autoantigens. All 3D structures of the human nervous system auto-antigens were obtained in PDB format from the AlphaFold database. The ID number of 3D structure of human nervous system autoantigens in the AlphaFold database is indicated in Table 1.

### Sequence alignment (definition of mimicry pentapeptides)

Our original program “Alignmentaj” [https://github.com/muslimb/MyProekt1] (Registered by Russian Federal Agency for Intellectual Property: Certificate # 2,023,617,186 of 6 April, 2023. https://www.elibrary.ru/item.asp?id=52295110) was used to identify mimicking pentapeptides between viral antigens namely EBV and HHV-6 and human CNS antigens namely MBP, MOG, MOBP, MAG, SYN1, SYN2, CLDN5, OCLN, TUBA1A and TUBB. As input, the program receives the amino acid sequences of human antigens and viral antigens in fasta format. The program divides the amino acid sequences of the virial antigens into pentapeptides (i.e., MSDEG, SDEGP, DEGPG, and so forth) and the program aligns the pentapeptides to the amino acid sequences of the human autoantigen. The output of the program shows existing mimicry pentapeptides between human autoantigens and microbial antigens.

### Analysis of Pentapeptide Mimicry Between EBV, HHV-6, and CNS Antigens for T and B Cell Immunogenicity

To investigate the immunogenic potential of the shared pentapeptides found in viral antigens (EBV and HHV-6) and different CNS proteins in stimulating T and B cells, we performed molecular mimicry analyses of these shared pentapeptides in relation to T and B cell activation. The Immune Epitope Database (IEDB) [40] was utilized to investigate the immunogenicity of mimicking pentapeptides of EBV and HHV-6 in relation to CNS antigens, focusing specifically on human immune cells, including T and B cells. The IEDB compiles experimental data regarding antibody and T cell epitopes that have been investigated in human subjects.

## Results

### The results of molecular mimicry between EBV and HHV-6 viral antigens and CNS proteins

#### EBV

In the present study, a total of 42 mimicry pentapeptides were identified between EBV and HHV-6 virial antigens and CNS antigens. As shown in **Table 2**, four mimicry pentapeptides were identified between EBNA1 and SYN1, while the same CNS proteins (SYN1) showed 3 mimicry pentapeptide with and EBNA6 (see **Figure 1A**). This CNS protein also showed 2 mimicry pentapeptide with LMP1 and EA-D viral antigens (see Figure 1A). Furthermore, only 1 mimicry pentapeptide was observed between SYN1 and EBNA2 and LMP2A/LMP2B as shown in Figure 1A. Myelin proteins, namely MBP (Figure 1B) and MAG (Figure 1C), showed two mimicry pentapeptides with EBNA1, while MAG showed one mimicry pentapeptide with EBNA2 and gE (see Figure 1C). MBP showed one single mimicry pentapeptide with EBNA6 and gE as presented in Figure 1B. Similarly, as displayed in Figure 1D, MOG showed one single mimicry pentapeptide with EBNA1, LMP1, BLLF1, and EBNA6.

**Figure 1.**
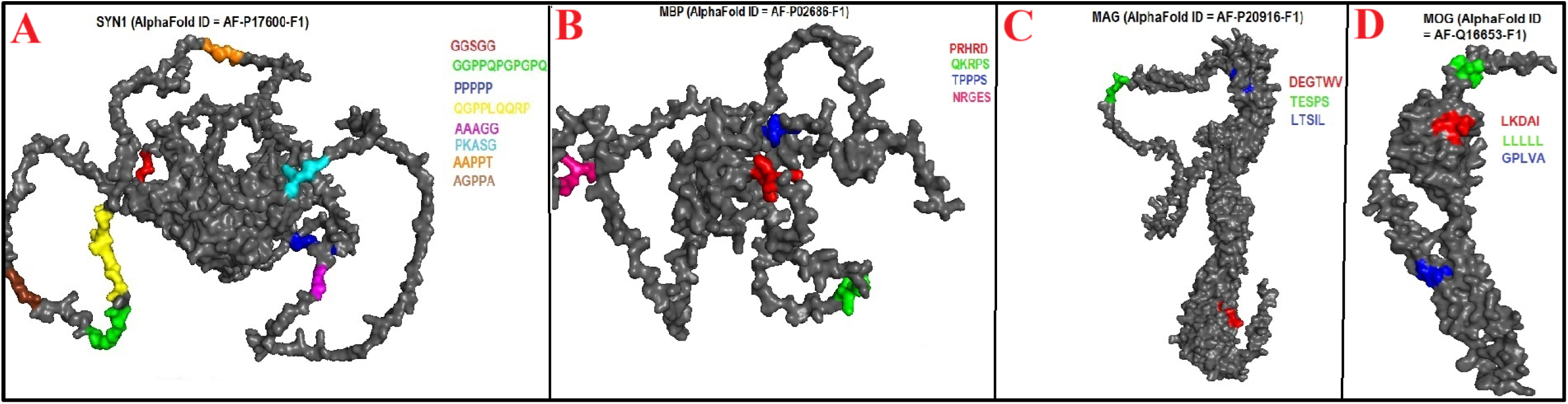
**(A-D)**. Location of Epstein–Barr virus (EBV) mimicking pentapeptides in 3D structures of syanpsin 1 (SYN1), myelin basic protein (MBP), myelin-associated glycoprotein (MAG), myelin oligodendrocyte glycoprotein (MOG).

**Table 2.**
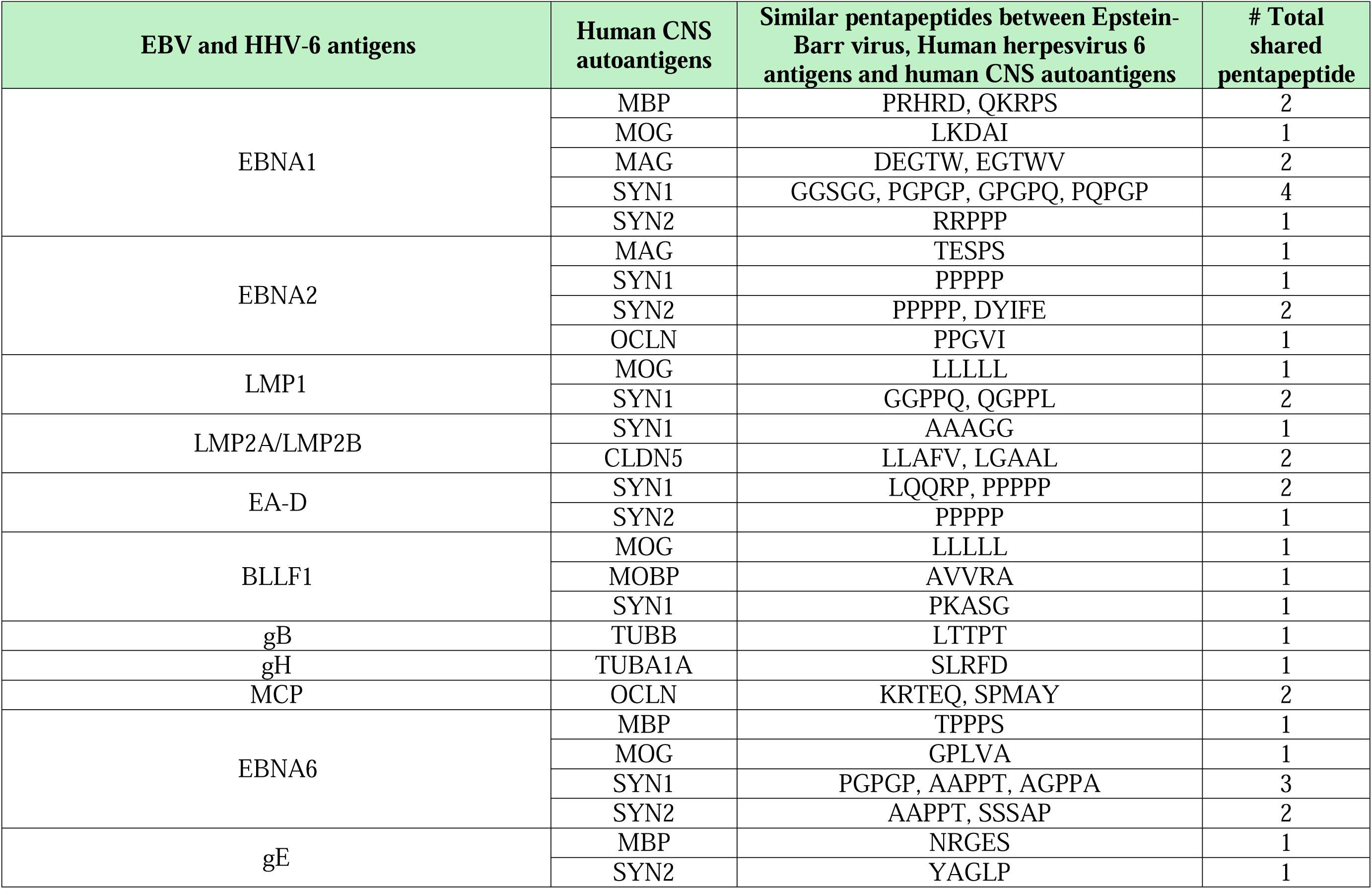
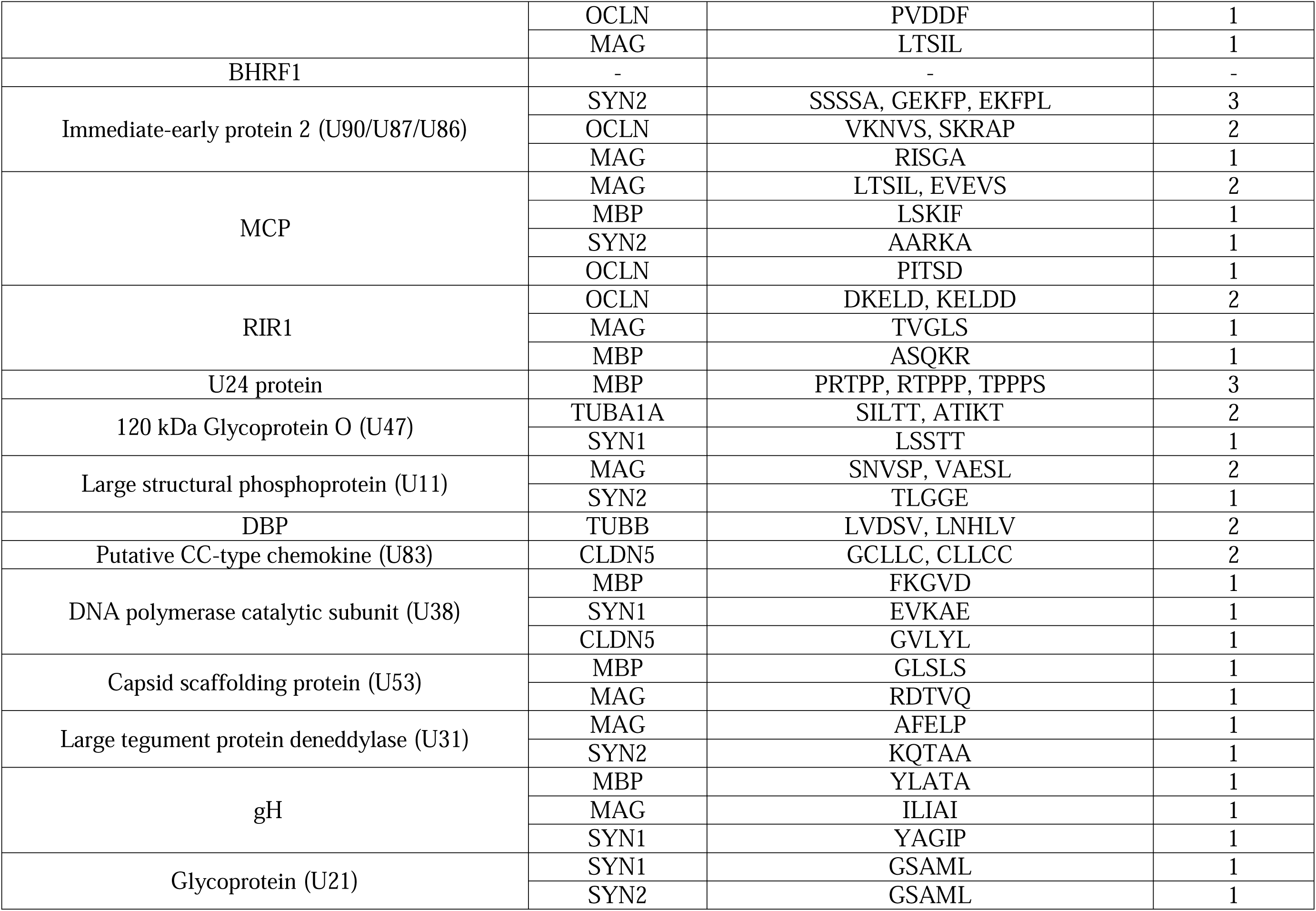
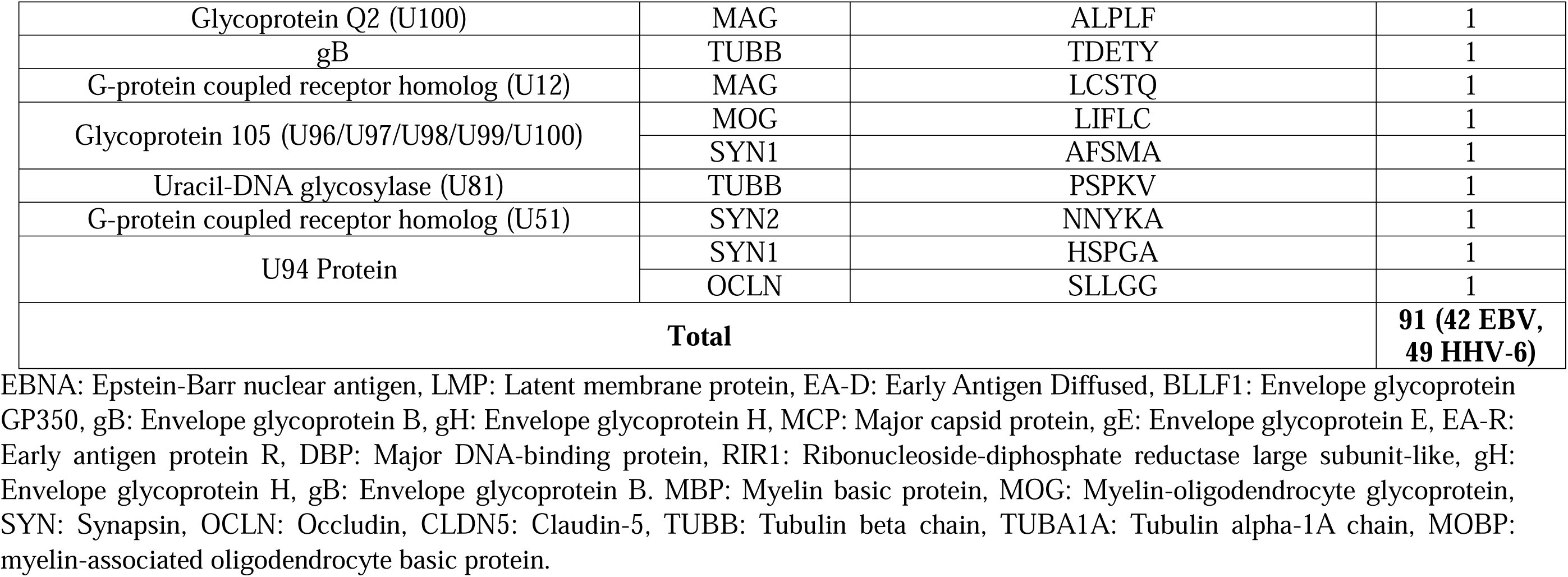
Results of molecular mimicry between Epstein–Barr virus (EBV), Human herpesvirus 6 (HHV-6) and central nervous system (CNS) antigens.

**Figure 2A** showed that SYN2 shared two mimicry pentapeptides with EBNA2 and EBNA6, and one single mimicry pentapeptide with EBNA1, EA-D, and gE. OCLN (Figure 2B) showed two mimicry pentapeptides with MCP and single one with EBNA2 and gE. CLDN5 (Figure 2C), MOBP (**Figure 3A**), TUBB (Figure 3B), and TUBA1A (Figure 3C) shared single mimicry pentapeptides with EBNA2, BLLF1, gB, and gH respectively.

**Figure 2.**
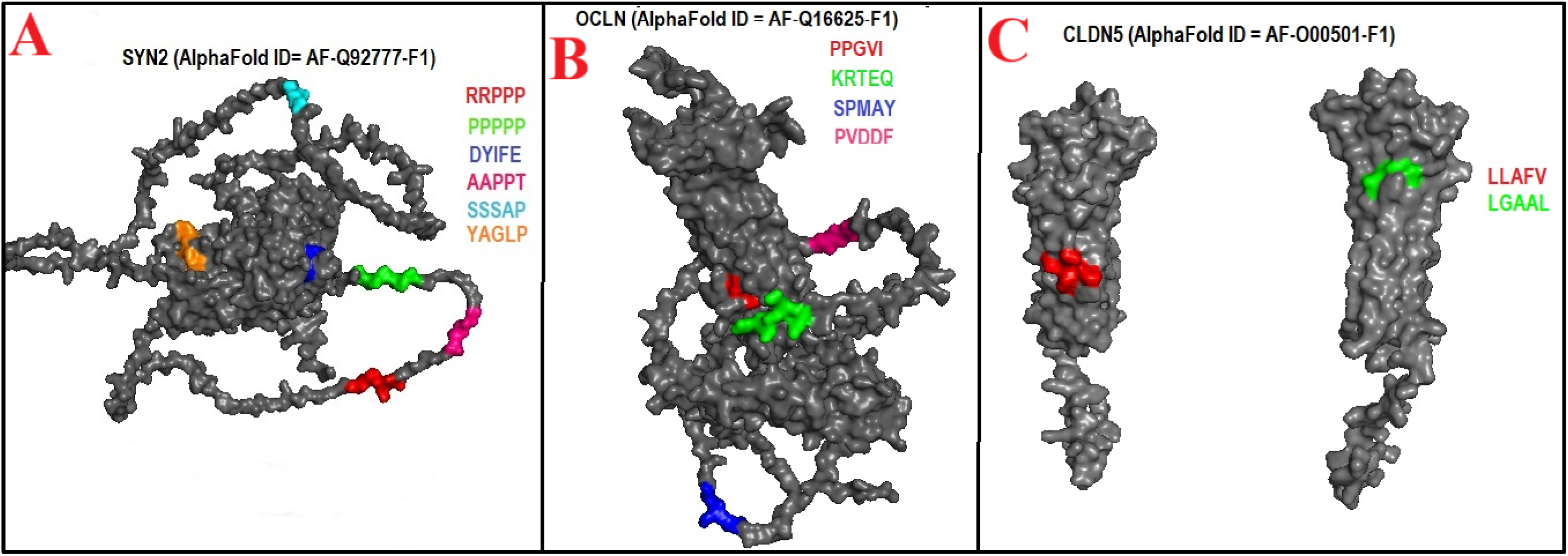
**(A-C)**. Location of Epstein–Barr virus (EBV) mimicking pentapeptides in 3D structures of synapsin (SYN2), occludin

**Figure 3.**
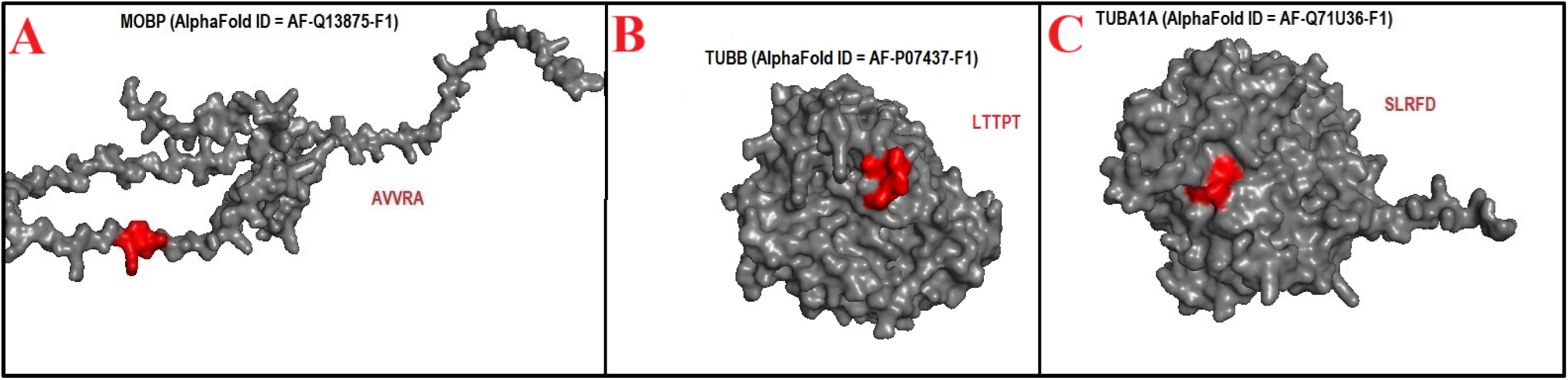
**(A-C)**. Location of Epstein–Barr virus (EBV) mimicking pentapeptides in 3D structures of myelin associated oligodendrocyte basic protein, tubulin beta (TUBB), tubulin alpha (TUBA1A).

#### HHV-6

The present study identified 54 mimicry pentapeptides shared between HHV-6 viral antigens and CNS antigens. As shown in Table 2 and **Figure 4A**, MBP exhibited three mimicry pentapeptides with the U24 protein and one with each of the MCP, RIR1, U38, U53, and gH. Likewise, Table 2 and Figure 4B indicate that SYN2 displayed three mimicry pentapeptides with immediate-early protein 2 and one with each of MCP, U11, U31, U21, and U51. Meanwhile, Table 2 and Figure 4C illustrate that MAG contained two mimicry pentapeptides with MCP and U11 alongside one with each of immediate-early protein 2, RIR1, U53, U31, gH, U100, and U12.

**Figure 4.**
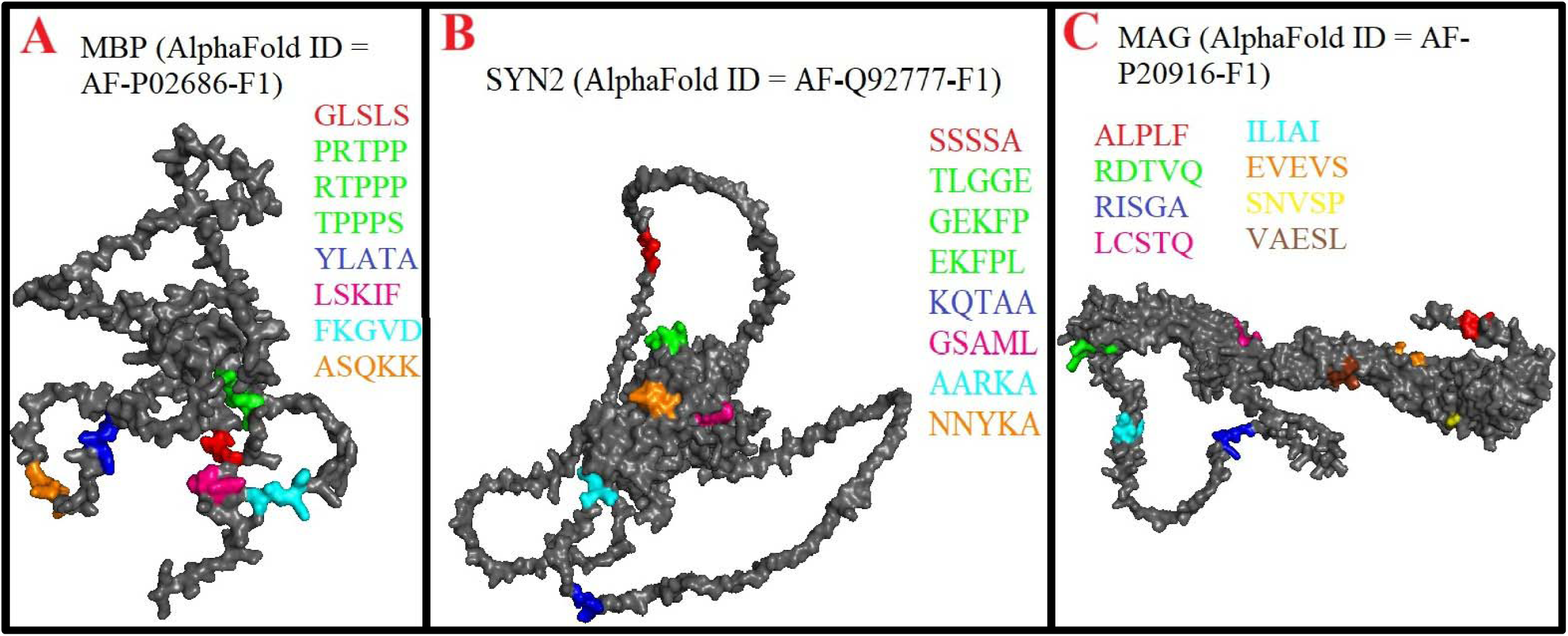
**(A-C)**. Location of Human herpesvirus 6 (HHV-6) mimicking pentapeptides in 3D structures of myelin basic protein (MBP), synapsin 2 (SYN2), myelin-associated glycoprotein (MAG).

Similarly, Table 2 and Figure 5A reveal that OCLN shared two mimicry pentapeptides with immediate-early protein 2 and RIR1, as well as one with MCP and U94 protein. Table 2 and Figure 5B demonstrate that SYN1 exhibited six mimicry pentapeptides, including one with each of the 120 kDa U47, U38, gH, U21, glycoprotein 105, and U94 protein. In addition, Table 2 and Figure 5C show that TUBB contained four mimicry pentapeptides, two with DBP and one each with gB and U81. Moreover, Table 2 and Figure 6A highlight that CLDN5 displayed two mimicry pentapeptides with the U83 and one with the U38. As depicted in Table 2 and Figure 6B, MOG exhibited one mimicry pentapeptide with glycoprotein 105, while TUBA1A contained one mimicry pentapeptide with the U47.

**Figure 5.**
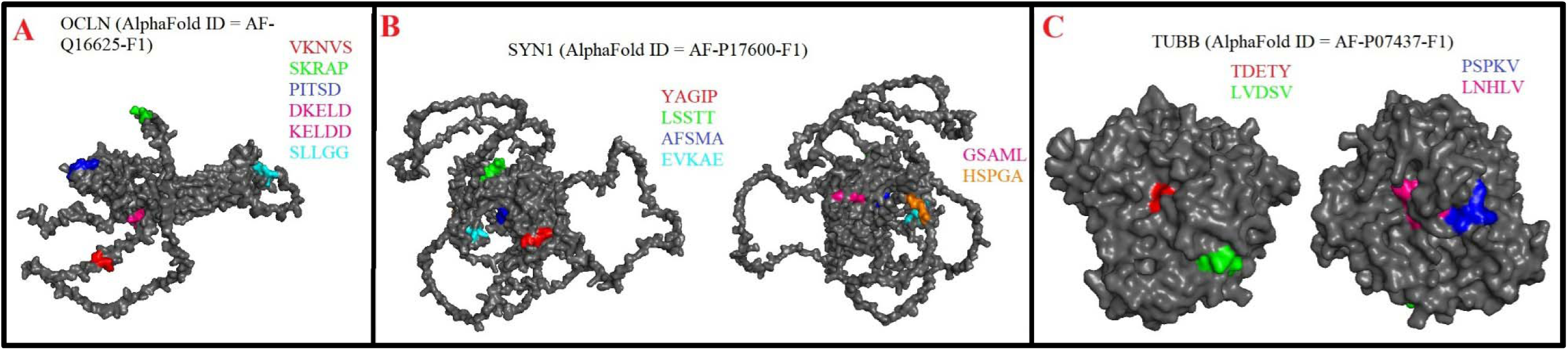
**(A-C)**. Location of Human herpesvirus 6 (HHV-6) mimicking pentapeptides in 3D structures of occludin (OCLN), synapsin 1 (SYN1), tubulin beta (TUBB).

**Figure 6.**
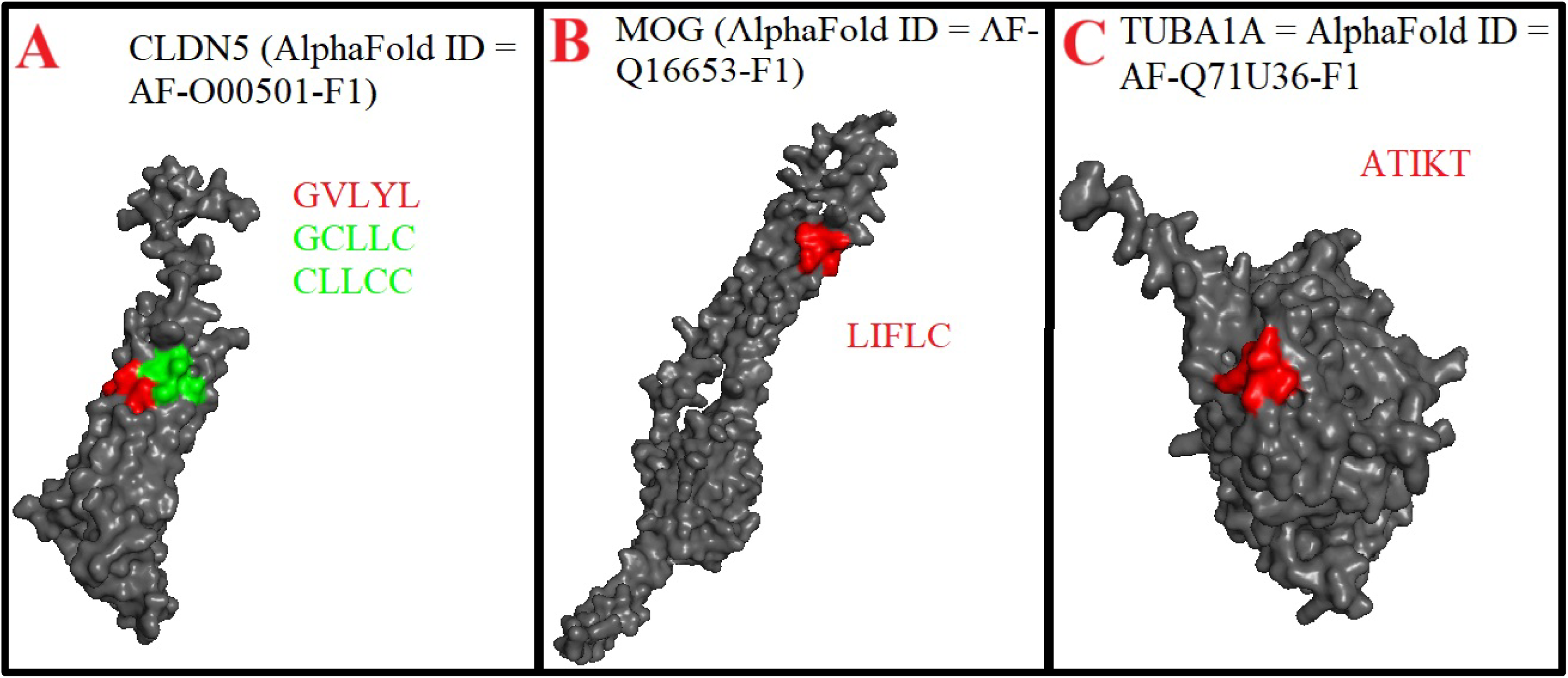
**(A-C)**. Location of Human herpesvirus 6 (HHV-6) mimicking pentapeptides in 3D structures of claudin 5 (CLDN5), myelin oligodendrocyte glycoprotein (MOG), tubulin alpha (TUBA1A).

### The immunogenic potentials of shared pentapeptides to trigger T and B cells

The immunoreactive epitopes of EBV and HHV-6, which share pentapeptide sequences with CNS proteins, are summarized in **Table 3**. The findings demonstrate that the homologous pentapeptides present in EBV nuclear antigens (EBNA1, EBNA2, and EBNA6), latent membrane proteins (LMP1 and LMP2), early antigen D (EA-D), and BLLF1 exhibit structural similarity with CNS autoantigens. These viral antigens can serve as immunoreactive epitopes targeted by both human T and B cells. Notably, all examined viral antigens, except for EBNA2, also share similar immunogenic pentapeptide sequences with SYN1. However, all viral antigens demonstrated similar immunogenic pentapeptides with other CNS proteins such as SYN2 and several key myelin-associated proteins, including MBP, MOG, and MAG, as well as the less common MOBP and CLDN5.

**Table 3.**
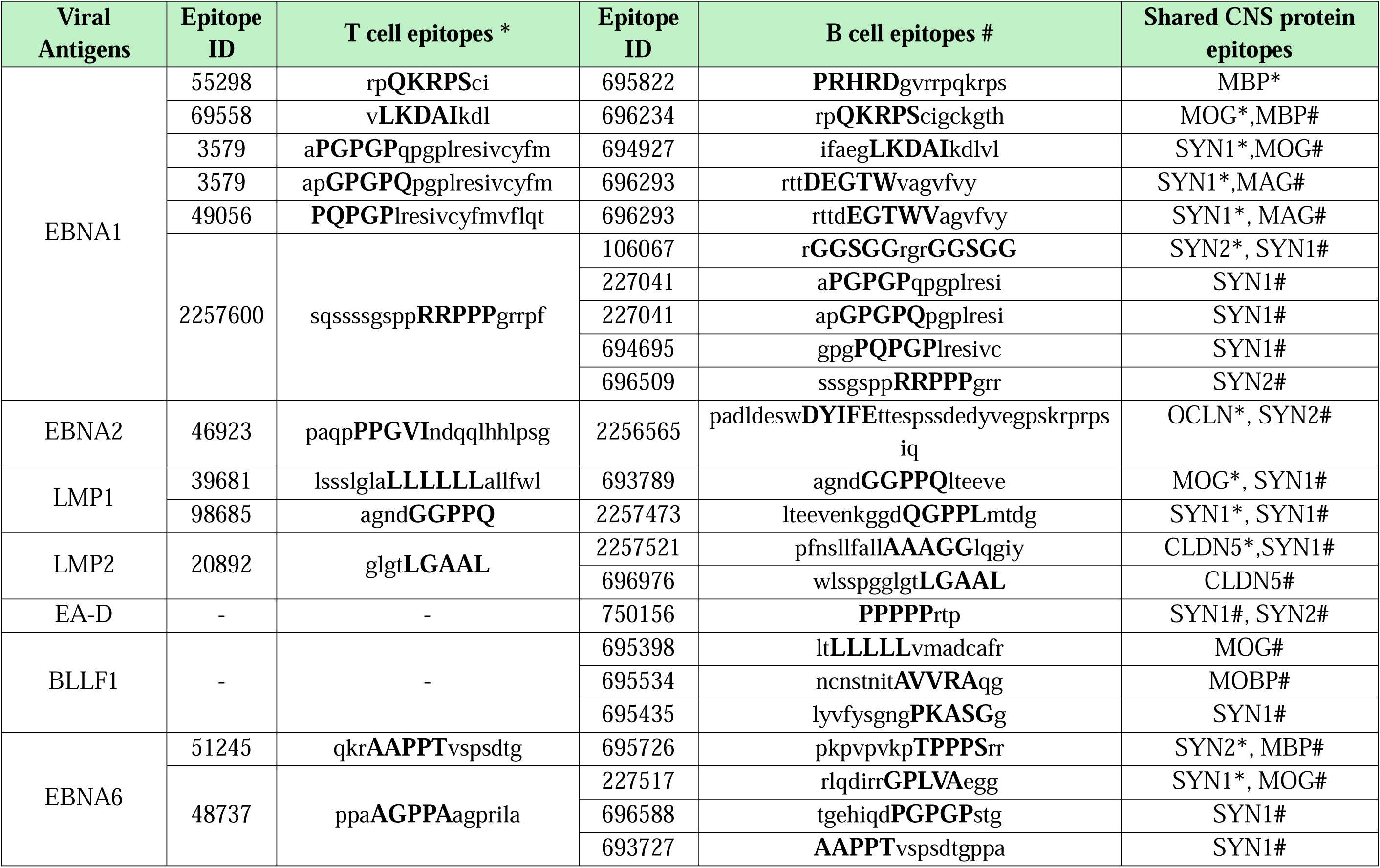
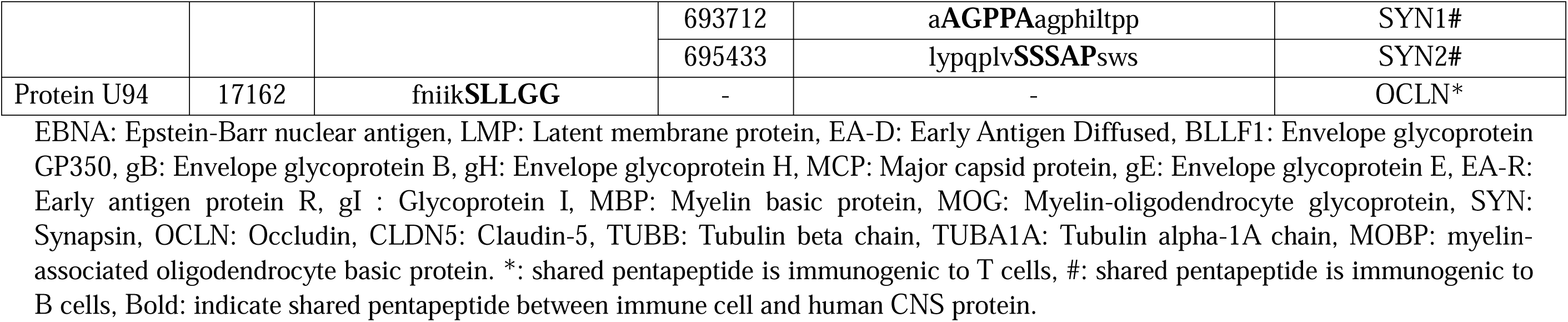
Immunoreactive Epstein–Barr virus (EBV), Human herpesvirus 6 (HHV-6)-derived epitopes containing shared pentapeptides with central nervous system (CNS) proteins. Shared pentapeptides are shown with bold.

## Discussion

### Shared pentapeptides between EBV and HHV-6 antigens and host cell CNS epitopes

The first key finding of the present study identified molecular mimicry between 12 EBV antigens (EBNA1, EBNA2, EBNA6, LMP1, LMP2A/LMP2B, EA-D, EA-R, gB, gH, MCP, gE and BLLF1) and 20 HHV-6 antigens (immediate-early protein 2, MCP, RIR1, U24, U47, U11, DBP, U83, U38, U53, U31, gH, U21, U100, gB, U12, glycoprotein 105, U81, U51, and U94) with 10 critical CNS proteins. These include myelin proteins (MBP, MOG, MAG, and MOBP), BBB tight junction proteins (OCLN and CLDN5), cytoskeletal proteins (TUBB, and TUBA1A), and synaptic proteins (SYN1 and SYN2). This mimicry suggests that immune responses to viral antigens may cross-react with structurally similar CNS self-epitopes, triggering immune-mediated CNS damage and contributing to autoimmunity. Unlike previous studies that focused on individual proteins, such as EBNA1 mimicry with GlialCAM [28], our study uniquely identifies mimicry with an expanded range of CNS proteins, including SYN1 and SYN2, and BBB-specific proteins OCLN and CLDN5. This enhances the comprehension of the range and variety of viral-induced autoimmune targets, especially in conditions characterized by impaired BBB and synaptic damage.

The immune mechanisms underlying this mimicry involve cross-reactive T cells and cross-reacting autoantibodies that damage host cell CNS epitopes and compromise the integrity of the BBB, facilitating the infiltration of peripheral immune cells into the CNS [3]. Such disruptions are hallmark features of CNS autoimmune diseases, particularly MS [3, 41] and are also prevalent in depression and CFS/ME (see Introduction).

Molecular mimicry likely functions via antigen presentation by MHC-II molecules, wherein viral shared motifs with host cell epitopes are displayed to autoreactive T cells, resulting in pathogenic immune activation [1, 8]. The subsequent upregulation of proinflammatory cytokines, including interferon-γ and tumor necrosis factor-α, exacerbate neuroinflammation and contributes to demyelination [42, 43]. EBV reactivation has been shown to expand autoreactive B and T cell populations, amplifying neuroinflammation and contributing to neuronal damage [44]. Similarly, HHV-6 latency within glial cells may induce proinflammatory cytokine release upon reactivation, further perpetuating neural injury [45]. These findings suggest that repeated viral reactivation or chronic exposure sustains inflammatory pathways and autoimmunity, offering a plausible mechanism for disease progression.

This mimicry has broader implications beyond MS, depression and CFS/ME. Research indicates that EBV and HHV-6 may play a role in various autoimmune and neurodegenerative diseases, including neuromyelitis optica, autoimmune encephalitis, and possibly Alzheimer’s disease, wherein viral reactivation has been linked to amyloid-β pathology [46–48]. The associations highlight the systemic influence of herpesviruses in neuroinflammatory and neurodegenerative disorders. Comparative analyses of other viral infections demonstrate unique mechanisms of molecular mimicry in CNS autoimmunity. For example, SARS-CoV-2 nucleocapsid protein shares homology with MS-associated proteins, such as PLP and neurofilament light polypeptide, implicating molecular mimicry in COVID-19-related neuroinflammation [49]. In contrast, EBV and HHV-6 exhibit mimicry with distinct CNS targets, including synaptic and cytoskeletal proteins, suggesting alternative pathways in autoimmune pathogenesis. The observed differences highlight the need for pathogen-specific strategies in therapeutic development, focusing on the unique mimicry profiles of individual viruses.

The identification of viral antigens that mimic key host cell CNS proteins has substantial therapeutic implications. Neutralizing monoclonal antibodies targeting viral antigens, such as EBNA1, could mitigate cross-reactive immune responses while preserving host tissue integrity [50]. Antiviral therapies that suppress EBV and HHV-6 reactivation could provide dual benefits by reducing viral load and dampening inflammation. Deleting EBV-infected cells while preserving healthy tissues are under investigation and may offer a promising strategy to reduce autoimmune activation without extensive immunosuppression [51]. Additionally, the design of vaccines that exclude mimicry-prone epitopes could prevent vaccine-induced autoimmunity while enhancing protective immunity [51].

The concept of molecular mimicry is not confined to viruses but extends to bacterial pathogens, reinforcing their role as a universal mechanism in autoimmune diseases. For instance, *Campylobacter jejuni* lipooligosaccharides mimic human gangliosides, triggering Guillain-Barré syndrome through an autoimmune attack on peripheral nerves [52]. Likewise, antigens from *Acanthamoeba castellanii* mimic myelin proteins, such as MBP and PLP, inducing experimental autoimmune encephalomyelitis (EAE), a murine model of MS [53]. Future research must validate these findings in animal models of neuropsychiatric disease, employ peptide-based methods to confirm cross-reactivity, and evaluate their clinical significance in larger cohorts of patients with CNS disorders and autoimmune disorders. These efforts may establish molecular mimicry as a key mechanism connecting viral infections to autoimmune pathogenesis and inform the development of precision-targeted interventions.

### T and B cells immunogenicity of the shared pentapeptide motifs

The second major finding of this study indicates that shared pentapeptides between EBV/HHV-6 antigens and human CNS proteins may elicit potent T- and B-cell-mediated immune responses. The SYN1 and SYN2 self-epitopes, which demonstrate shared pentapeptides homology with various EBV antigens, were identified as significant stimulators of immune activation among the shared proteins. This is consistent with recent findings that emphasize the significance of synaptic proteins in autoimmune diseases characterized by neuroinflammation and synaptic dysfunction as key features of the disease pathology [54].

Previous research indicates that MS and the affective and CFS/ME symptoms due to MS are linked to autoimmune responses involving IgA, IgG, and IgM antibodies that target critical myelin proteins, including MBP, MOG, and MAG [12]. This study extends existing evidence by proposing that autoimmune responses may be partially influenced by increased immune reactivity to viral antigens, such as EBNA and deoxyuridine-triphosphate nucleotidohydrolases from EBV and HHV-6 [12]. In Long COVID, antibodies directed against HHV-6 epitopes correlate with elevated IgA and IgG titters against zonulin, a crucial protein for intestinal barrier integrity [14]. This cross-reactivity highlights the systemic effects of molecular mimicry and its possible involvement in sustaining neuroinflammatory and autoimmune mechanisms.

The immunogenicity of shared epitopes is further supported by Begum et al. (2022), who identified 13 T-cell epitopes shared between viral and human proteins, capable of binding promiscuously to human HLA class II alleles [55]. As mentioned earlier in this study, these epitopes likely trigger autoreactive T cells through mechanisms of molecular mimicry, which is well-documented in autoimmune diseases such as MS [3]. Human coronaviruses and other pathogens have been implicated in activating myelin-reactive T cells by presenting antigens that resemble CNS proteins [56, 57]. Similarly, in HAM/TSP, molecular mimicry has been shown to involve cross-reactive antibodies targeting HTLV-1-tax and hnRNP A1, a neuronal protein, contributing to CNS damage [58].

EBV encodes multiple proteins that may disrupt immune tolerance by mimicking host immune-regulatory genes. For instance, BCRF1, an IL-10 homolog, modulates cytokine production, while BZLF-1, a mimic of AP-1 and NF-κB, promotes proinflammatory pathways [59]. Moreover, exosomes released from EBV-infected B cells contain viral components capable of triggering systemic and CNS-specific immune responses, contributing to the development of MS [60]. These mechanisms are complemented by herpesviruses’ ability to reduce epitope diversity and enhance similarity to host peptides, thereby evading immune detection and facilitating chronic infection [61]. Such strategies allow these viruses to persist in the host while simultaneously driving autoimmune pathology through prolonged immune activation.

CD8 T cells recognize specific EBV antigens, such as EBNA1, presented via MHC class I molecules. For example, EBNA1 peptides restricted by HLA-B8, HLA-B3501, and HLA-Cw0303 elicit strong T-cell responses, although the glycine-alanine repeat (GAr) domain in EBNA1 inhibits proteasomal degradation, limiting peptide availability for presentation [62, 63]. Despite this, epitopes flanking the GAr domain remain accessible and are sufficient to activate CD8 T cells [62]. Likewise, CD4 T cells recognize EBNA1 peptides presented by MHC class II molecules, although the protein’s low expression often limits efficient presentation [64]. Nevertheless, specific epitopes, such as EYHQEGGPD, are effectively presented and induce robust CD4 T-cell responses, highlighting their potential as therapeutic targets in immunotherapy [64].

MBP-specific T cells can be activated by viral antigens, such as EBV LMP1, through shared epitopes, promoting epitope spreading and enhancing autoreactive B-cell responses [65]. This mechanism is critical in MS, where chronic viral reactivation redirects immune responses toward myelin proteins, amplifying neuroinflammation [66]. Similarly, MOG peptides bind efficiently to both MHC class I and II molecules, eliciting strong T-cell responses in inflammatory demyelinating diseases [67, 68].

Antigens from EBV and HHV-9, including dUTPase, activate the TLR2 and NFκB signaling pathways, thereby amplifying inflammatory responses that contribute to autoimmune activation [69]. Shared epitopes between EBNA1 and CNS proteins, such as alpha-crystallin B (CRYAB), facilitate cross-reactive T-cell responses that are implicated in MS and other neuroinflammatory disorders [70]. Additionally, herpesviruses exploit latency phases to express immune-evasive proteins that mimic host peptides, promote chronic infection and sustained immune activation [61, 71]. This dynamic interplay between immune evasion and mimicry establishes herpesviruses as significant contributors to autoimmune pathogenesis.

## Limitations

The in-silico approach, although effective for predicting molecular interactions and sequence homology, fails to capture the full complexity of host-pathogen interactions in vivo. The immune responses elicited by mimicry require validation in animal models or human clinical samples. A further limitation is the insufficient data regarding the temporal relationship between viral reactivation and the onset or progression of autoimmune neuroinflammatory diseases, necessitating longitudinal studies. The integration of advanced technologies, including single-cell RNA sequencing and high-resolution imaging, may enhance the understanding of the cellular and molecular mechanisms underlying mimicry-induced pathologies [72]. These approaches will enhance the evidence base and promote the application of mimicry-based findings in clinical practice. This study examined molecular mimicry only in HHV-6A, without assessing HHV-6B, which has distinct immune interactions. Future research should investigate HHV-6B to determine its potential role in immune cross-reactivity

## Conclusion

This study identifies a significant overlap of pentapeptides between EBV and HHV-6 antigens and key CNS proteins, underscoring a potential mechanism by which viral infections contribute to CNS autoimmunity through molecular mimicry. The structural similarity between these viral and host cell epitopes may elicit robust immunogenic responses from both T and B cells, suggesting that immune cross-reactivity plays a critical role in the pathogenesis of CNS diseases. These findings reinforce the hypothesis that persistent viral infections or reactivation events may drive autoimmunity by priming autoreactive lymphocytes against self-antigens, ultimately leading to neuroinflammatory damage. Given the implications of these interactions in neuropsychiatric disease, further investigations are warranted to delineate the precise molecular pathways involved and to explore targeted therapeutic strategies that could mitigate virus-driven autoimmune responses.

## Ethical approval and consent to participate

None.

## Availability of data and materials

MGN will reply to reasonable requests for the dataset used in the current study after all authors have fully utilized the data.

## Credit author’s contributions!

AFA and MGN carried out the current study’s design. The data was gathered by AFA, TS, MGN. Bioinformatics analysis was performed by MGN. AFA and MM wrote the first draft. All authors contributed to the editing of the work, and they have all given their consent for submission of the completed version.

## Acknowledgments

Not applicable.

